# Relationship between bacterial phylotype and specialized metabolite production in the culturable microbiome of two freshwater sponges

**DOI:** 10.1101/2021.08.26.457769

**Authors:** Chase M. Clark, Antonio Hernandez, Michael W. Mullowney, Jhewelle Fitz-Henley, Emma Li, Sean B. Romanowski, Roberto Pronzato, Renata Manconi, Brian T. Murphy

## Abstract

Microbial drug discovery programs rely heavily on accessing bacterial diversity from the environment to acquire new specialized metabolite (SM) lead compounds for the therapeutic pipeline. Therefore, knowledge of how certain bacterial taxa are distributed in nature, in addition to the degree of variation of SM production within those taxa, is critical to informing these front-end discovery efforts and making the overall sample collection and bacterial library creation process more efficient. In the current study we employed MALDI-TOF mass spectrometry and the bioinformatics pipeline IDBac to analyze diversity within phylotype groupings and SM profiles of hundreds of bacterial isolates from two *Eunapius fragilis* freshwater sponges, collected 1.5 km apart. We demonstrated that within two sponge samples of the same species, the culturable bacterial populations contained significant overlap in approximate genus-level phylotypes but mostly non-overlapping populations of isolates when grouped lower than the level of genus. Further, correlations between bacterial phylotype and SM production varied at the species level and below, suggesting SM distribution within bacterial taxa must be analyzed on a case-by-case basis. Our results suggest that two *E. fragilis* freshwater sponges collected in similar environments can exhibit large culturable diversity on a species-level scale, thus researchers should scrutinize the isolates with analyses that take both phylogeny and SM production into account in order to optimize the chemical space entering into a downstream bacterial library.

## Introduction

Marine sponges [1, 2] contribute significantly to known chemical space [3], accounting for nearly 30% of all specialized metabolites (SMs) reported from the marine environment [1, 4]. It is also increasingly apparent that microorganisms play an important role in shaping the chemical space of the sponge holobiont [5, 6]. However, sponges that live in freshwater environments remain relatively understudied [7–10], as do their microbiomes. In the Great Lakes and neighboring regions, freshwater sponge distribution and diversity has been documented since explorations occurring as early as the late 19^th^ to early 20^th^ century [11–15], as well as in recent efforts [16, 17]. While mostly ecological studies on freshwater sponges have documented the phospholipid, fatty acid, and sterol composition of whole-sponge extracts [18–25], there have been limited studies on the diversity of their bacterial communities [26–31] and the associated SM capacity within these communities.

Since 2015 our team has worked with citizen scientist SCUBA divers in the Great Lakes region of the United States to collect freshwater sponges as sources of bacterial isolates for our SM drug discovery pipeline. Historically, when samples are collected from the environment, differences in both the phylogenetic diversity and the SM capacity of the culturable microbial population are not fully considered when creating a bacterial library. However, this information is critical to determine the extent to which samples should be collected, particularly when operating under a limited research budget, as is often the case in academia [32]. Understanding the bacterial phylogenetic and SM variance between sponge samples would allow for a more structured design of collections and would make the overall sample collection and bacterial library creation process more efficient [33, 34].

With this in mind, two samples of the freshwater sponge species *Eunapius fragilis* (Leidy, 1851) (Porifera: Demospongiae: Spongillida: Spongillidae) were collected 1.5 km apart in the St. Lawrence River. The collected tissue was prepared using protocols described in the experimental section, and every distinguishable bacterial colony was isolated from nutrient media plates. We focused solely on the readily culturable population because this reflects the methodological constraints of many modern microbial drug discovery programs. Protein and SM mass spectra were collected using matrix assisted laser desorption/ionization time-of-flight mass spectrometry (MALDI-TOF MS) and analyzed using our previously developed open-source bioinformatics software IDBac to simultaneously assess the proteomic and metabolomic variation of the resulting isolates [29, 30]. This allowed us to investigate differences in culturable bacterial pseudo-phylogenetic and SM populations between two *E. fragilis* samples collected close in location and time. Specifically, we examined 1) the extent to which readily-culturable bacterial populations varied between the two sponges, and 2) patterns of SM variation within groups of closely related bacterial isolates.

## Results/Discussion

### Culturing the *Eunapius fragilis* microbiome

Two samples of *E. fragilis*, were collected from the St. Lawrence River, USA approximately 30 km from the outlet of Lake Ontario. These samples were collected 1.5 km apart, in both a similar time window and lifecycle, and are referenced herein as sponges “SCD18” and “SCD21”. Both sponge samples were processed simultaneously, using identical protocols. After thoroughly washing sponge tissue in sterilized freshwater, 108 microbial diversity plates were generated by grinding the tissue in sterile 20% glycerol and applying two pretreatment conditions (with and without heat treatment), three dilution series, and nine agar-based nutrient plate formulations. As expected, some of the resulting plates exhibited no discernible colonies or were overgrown with bacterial biofilms or cycloheximide-resistant fungus. For the remaining plates every distinguishable colony was then isolated, resulting in 522 isolates from sponge SCD18 and 329 isolates from sponge SCD21 (total of 851 isolates). To compare bacterial diversity equally between the two sponge samples, only isolates derived from matching sample plates were included in our analysis (e.g., if a nutrient agar diversity plate was contaminated with a fungus, the plate with matching conditions from the second sponge was removed from analysis as well). This resulted in a total of 692 bacterial isolates that were advanced to our MALDI-TOF MS pipeline, where protein and SM spectra were acquired for 3 biological replicates of each isolate.

### Documenting the pseudo-phylogenetic bacterial diversity within *Eunapius fragilis*

MALDI-TOF MS protein spectra (3000 to 15000 Daltons) consist primarily of ions of intact ribosomal, cell structure, and regulatory proteins [35, 36] and are often used to group microorganisms at the genus, species, and sub-species taxonomic levels (a few examples [37–43]). This technology has been applied extensively in clinical settings [43, 44]. However, the widespread adoption of MALDI-TOF MS for the study of environmental bacteria has lagged behind clinical applications [45]. This is primarily due to the lack of reference spectra needed to identify unknown isolates and a scarcity of MALDI-TOF MS analysis tools that are freely available to the public [38, 39, 46–48]. We previously designed IDBac as an open-source bioinformatic tool to first cluster bacterial isolates by protein spectrum similarity, followed by subgrouping them based on feature overlap within corresponding SM spectra (<2000 Daltons) [49, 50]. A unique advantage of IDBac is that it combines proteomic and metabolomic data into a semi-automated pipeline that facilitates the assessment of microbial diversity beyond the scope of typical MALDI-TOF MS pseudo-phylogenetic analyses.

In order to assess the taxonomic diversity and overlap between all 851 bacterial isolates from both sponge samples, IDBac was used to cluster isolates into a dendrogram (Fig. S1) by measuring the cosine similarity of MALDI-TOF MS protein spectra [39] and clustering with Ward’s method [51, 52]. The resulting groupings were evaluated for accuracy by subsampling isolates across the dendrogram and using 16S rRNA gene sequencing analysis to obtain genus-level identifications. Additional genus-level assignments were made by seeding MALDI-TOF MS protein spectra of previously identified isolates [34] into the dataset. Taxonomic assignments made by MALDI-TOF MS and 16S rRNA analyses complemented one another (Figs. S2 and S3).

In addition to confirming accurate clustering, these complementary techniques revealed that isolates spanned at least four phyla commonly associated with sponges: Proteobacteria, Actinobacteria, Bacteroidetes, and Firmicutes (Fig. S4). These phyla are responsible for greater than half of known microbially-produced SMs to date [32, 53]. In total, 11 genera were identified in the dendrogram, though more were cultivated and remain unidentified (Figs. S1-S4). This initial organization of sponge isolates into a dendrogram allowed for more in-depth analyses, including comparison of recovered bacterial communities between the two sponges, and the overlap of SM features within phylotypes.

### Measuring variation between freshwater sponge culturable bacterial populations using MALDI-TOF MS

Unsupervised machine learning techniques such as hierarchical clustering are often used to group MALDI-TOF MS protein spectra of bacteria [38, 39, 49]. One method to create discrete groups from hierarchical clustering (visualized as a dendrogram) is to “cut” across the axis representing distance (Fig. 2a). However, a notable limitation in hierarchical clustering of MALDI-TOF MS spectra is its inability to consistently link last common ancestors at higher taxonomic ranks such as Family and above. This limitation extends in some cases to highly diverse genera such as *Streptomyces* and *Bacillus*. This is attributed to the widespread use of similarity measures (by all current intact-cell MALDI-TOF MS analyses, including IDBac) that cannot account for biologically based chemical modifications, however minor, which result in a mass shift. Because MALDI-TOF MS spectrum similarity does not have a linear correlation with 16S rRNA similarity [39], inferred pseudo-phylogenetic relationships made using MALDI-TOF MS spectra are most accurate when comparing isolates at the species to subspecies levels [38, 39, 49, 54]. In other words, MALDI-TOF MS dendrograms can accurately group isolates of the same species and subspecies, but often will not cluster higher taxonomic ranks together.

Groupings from hierarchical clustering of MALDI-TOF spectra are more accurate with an increasing number of isolates. For this reason, we performed initial clustering using all 851 isolates (Fig. S1), but then pruned 159 isolates from non-matched diversity plates from the full dendrogram (Fig. 1). This left a total of 692 recovered bacterial isolates that were subjected to comparative analyses.

**Figure 1.**
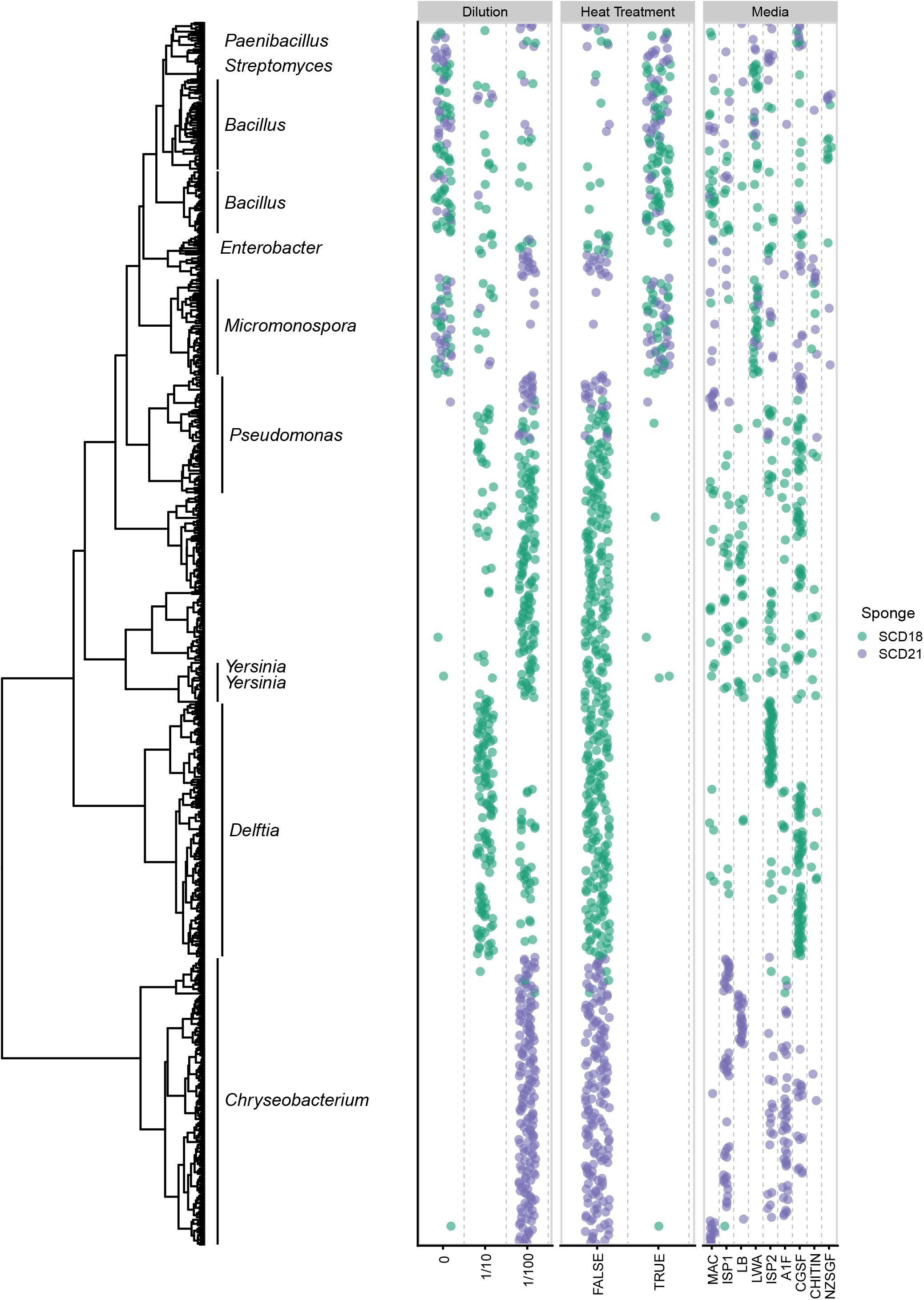
IDBac dendrogram of culturable bacterial population from freshwater sponges *Eunapius fragilis* SCD18 and SCD21. Dendrogram of 692 sponge-derived bacterial isolates, grouped by MALDI-TOF MS protein spectra similarity. The occurrence of bacterial isolates as a function of growth condition was mapped onto the dendrogram and color coded by source sponge.

The pruned dendrogram was cut 160 times at height intervals of 0.05, and for every cut, the composition of the resulting pseudo-phylogenetic groups (phylotypes) was tallied, whether groups contained isolates from SCD18, SCD21, or both; Fig. 2. Between cut heights ranging from 0-2 (roughly species-level and below), most groups consisted of isolates from either SCD18 or SCD21, but not both. When the dendrogram was cut at a height greater than 2.5 (roughly genus level and above), groups largely consisted of isolates from both sponges, except for the *Delftia* group which consisted of isolates only from SCD18. This suggests that at the bacterial species level, most phylotypes were only present in either one sponge or the other. This conclusion was supported by observing similar results from hierarchical clustering using a different peak binning algorithm and distance measure (Fig. S5), and additional analysis by k-means clustering (Fig. S6).

**Figure 2.**
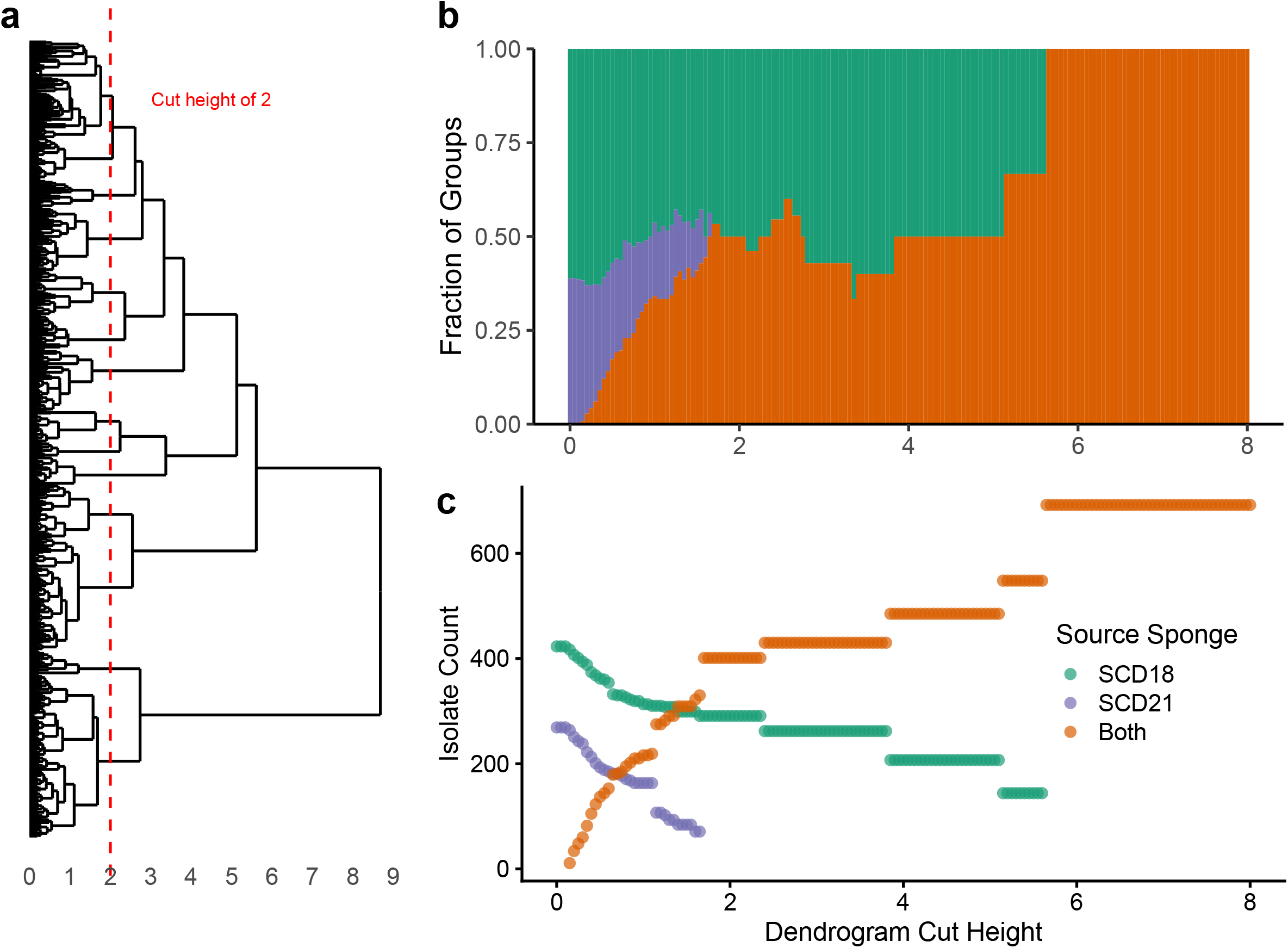
When cutting the dendrogram at a height that approximates species-level groupings and lower (a), most groups of isolates occurred in either one sponge or the other, not both, indicating a rich culturable species-level variation in the two *E. fragilis* samples collected 1.5 km apart (b). Because the number of observed groups decreases significantly as the dendrogram cut height increases, raw isolate counts were evaluated within groups for every cut (c). In the case of the depiction of groups (b) belonging to “Both” (in orange) is identical to the Jaccard index.

Many of the clusters identified at the genus level in Fig. 2 could be defined by linkage heights between 2 and 3, while much of the variation between sponges occurred at or below a height of 2. In order to confirm that the different populations were not simply the result of experimental MS artifacts, a phylogenetic tree was created with SILVA ACT [55] from 16S rRNA gene sequences of isolates and closely related reference strains (Fig. S7). This revealed that identified genera were not composed of a single species. It also revealed that cuts of the protein spectra dendrogram at a height of approximately 2 and below (where variation between sponges was highest) represented true species differentiation, not simply MS artifacts.

These results are likely not representative of the relative abundances of in-situ bacterial populations [56], rather they represent the populations of culturable bacteria that are easily accessible to researchers who engage in microbial drug discovery efforts. These results suggest that although the readily recoverable bacterial populations from two *E. fragilis* samples collected in similar location and time windows shared many of the same genera, they also varied significantly at the species, and possibly sub-species level. Since phylogenetic diversification is correlated with differential SM production capacity, this information has great potential to drive future SM discovery efforts. With this in mind, it was important to analyze whether isolates within similar phylotypes harbored overlapping SM mass features.

### The relationship between phylotype and chemotype in cultured *Eunapius fragilis* bacteria

Since the re-isolation of known SM scaffolds from redundant entries in a microbial library is a major concern for researchers engaging in microbial drug discovery, it was necessary to document the degree of SM overlap within dendrogram groupings. When determining the overlap of SMs between cultured bacterial isolates, it is important to consider that many bacterial SMs are encoded by co-localized genes within biosynthetic gene clusters (BGCs). Within genera, the loss and/or functional replacement of BGCs [57, 58] are critical forces that drive SM diversity within and between bacterial communities. This, in addition to transcriptional and translational regulation of BGCs, means that SMs may be observed frequently across multiple species in a given genus or may be specific to a single species. To determine the intra-genus SM variation among our freshwater sponge bacterial isolates, we generated metabolite association networks (MANs) from dendrogram groupings, an embedded function within IDBac.

MANs were used to analyze MALDI-TOF SM spectra by linking isolates with shared mass features (Fig. 3) [49]. Larger colored circles represent individual bacterial isolates, while smaller gray circles are mass features in the spectra, which commonly represent SMs [49]. For this analysis a subset of six genus-level phylotypes were chosen based on their historic precedent for producing SMs (*Streptomyces, Bacillus, Micromonospora, Pseudomonas;* these account for over half of documented bacterial SMs to date) or due to the abundance of recovered isolates from sponge samples (*Chryseobacterium* and *Delftia*; these represented 45% of the cultured isolates obtained from both sponges) [40]. Representative isolates from these six genera were identified either by matching MALDI-TOF MS spectra to spectra from characterized isolates in our in-house library and/or 16S rRNA gene sequence analysis (Fig. S2-S3).

**Figure 3.**
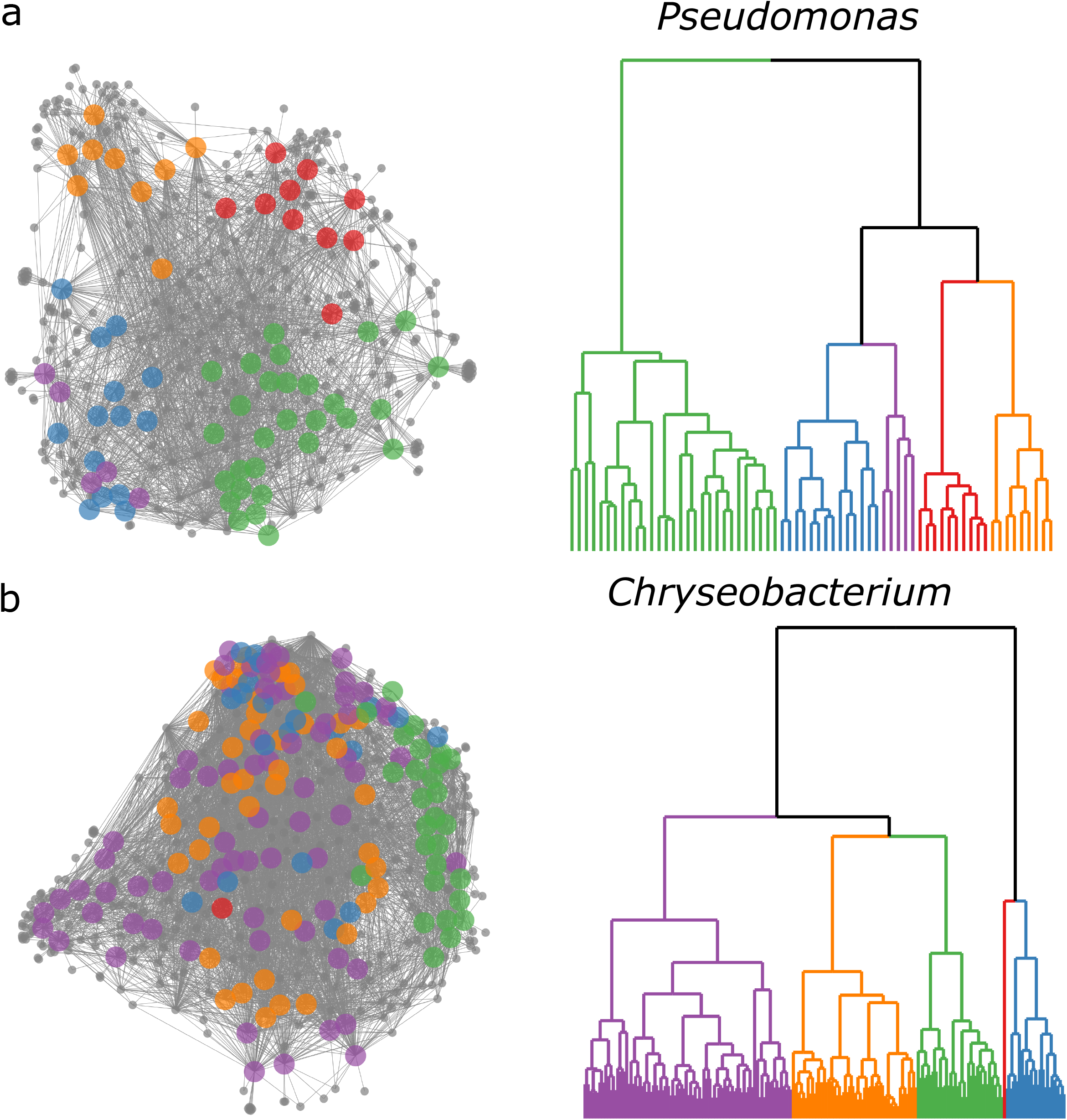
Metabolite Association Networks (MANs) of *Pseudomonas* and *Chryseobacterium* dendrogram groupings. Examples of MANs that show high (a) and low (b) correlation with protein spectra dendrogram groupings. Dendrograms are approximately the same height 2.36 (a) and 2.73 (b); and both were split into 5 groups. MANs are colored according to subgroupings in their corresponding pseudo-phylogenetic dendrograms. *Pseudomonas* isolates showed SM variation that correlated with protein-based dendrogram groupings (a). Conversely, *Chryseobacterium* showed only marginal correlation (b).

Across the dataset, the largest observed influence on SM spectrum similarity was protein spectrum similarity, which is a proxy for phylogenetic relatedness. Positive Pearson correlation coefficients between protein and SM spectra for the six groups mentioned are displayed in Fig. S8, in addition to pairwise comparisons between all isolates in Figs. S9 and S10. This phenomenon is clearly demonstrated in the MAN of the *Pseudomonas* dendrogram grouping (Fig. 3a), which contained sub-groups of isolates that exhibited differential SM production patterns and correlated directly to breaks in the dendrogram (Fig. 3b). This correlation aligns well with other recent studies [59–64] and highlights that species-level phylogenetic identity can be an effective guide for predicting SM diversity.

However, while some groupings showed high correlation of protein and SM spectra similarities at the species level, exceptions did exist. For example, unlike with *Pseudomonas*, isolates within the *Chryseobacterium* MAN (Fig. 3b) showed significant SM overlap that was less correlated to protein spectrum similarity (Fig. 3b and S8). Possible explanations for this include 1) SM BGCs are relatively conserved within the *Chryseobacterium* genus and/or this set of isolates (recent preliminary evidence supports this, though extensive investigation is needed to confirm this [65]); 2) *Chryseobacterium* isolates exhibit fewer expressed SMs under these cultivation conditions compared to the *Pseudomonas* isolates, and as a result provide fewer opportunities to observe differentiation; 3) different resolution of the MALDI-TOF MS protein pseudo-phylogeny was achieved between *Chryseobacterium* and *Pseudomonas* isolates. However, MALDI-TOF analysis, which specializes in species/sub-species differentiation, did delineate several distinct groups of *Chryseobacterium* isolates below the level of genus (Fig. S12–S14). Observing varying relationship of phylotype to chemotype (Figs. S8-S11a) highlights the need to resolve isolates using orthogonal methods, such as protein *and* SM spectra similarity.

Lastly, SM variation as a function of source sponge was analyzed within the six genera (Fig. S11b). When analyzing isolates within each genus-level dendrogram grouping, MANs did not exhibit any noticeable partition of SM mass features according to their source sponge, indicating that if a bacterial species occurred in both sponges, it was likely to produce similar suites of SMs. This indicates that in the dataset analyzed, phylogeny was a better prediction of SM production similarity rather than sample source.

The positive correlation of phylogeny with SM production, though generally true, is not sufficient to fully annotate the extent to which SMs are distributed within bacterial taxa. This principle was recently demonstrated on large scale analyses of the human gut microbiome [66]. More complex correlations between phylotype and SM production that align with this principle are exhibited in each of the MANs of *Micromonospora, Streptomyces, Bacillus*, and *Delftia* groups (Fig. S11a). To gain greater insight of SM variation within a large bacterial library, both phylogenetic and metabolomic analyses should be undertaken. A linear-mode-only MALDI-TOF MS instrument (limited to acquiring protein spectra) is sufficient to provide species to subspecies resolution of a bacterial isolate collection, and a dendrogram of those isolates may be a strong indicator of SM variation. However, to observe sub-genus SM variation, linear *and* reflectron (SM analysis) mode MALDI-TOF MS is required.

A major obstacle to current microbial-based drug discovery campaigns is the accumulation of isolates in a library that contain highly overlapping SM populations. Taxonomic and SM redundancy greatly reduce drug discovery efficiency, particularly in resource-limited academic laboratories. Our findings show that morphological-based colony isolation tends to be reductionist, as it does not take into consideration the complexities of vertical and horizontal transfer of SM BGCs. For example, SM populations in a drug discovery library could be highly redundant if researchers incorporated multiple isolates from the sponge *Chryseobacterium* group in Fig. 3b. Alternatively, unique SM populations could be overlooked in the case of the *Pseudomonas* group, where although the gross SM mass features appear largely homogeneous, many of the MAN subgroupings contain unique mass features that may correspond to a SM that is highly specific to a given bacterial strain, and may hold a unique ecological purpose and biological activity. These complexities highlight the need to consider both phylogeny and SM production on a case-by-case basis.

### Conclusions

The current study revealed that MALDI-TOF MS and IDBac are valuable tools for post-hoc analyses of culturable bacterial isolates. We demonstrated that within two sponge samples of the same species, the culturable bacterial populations contained significant overlap in approximate genus-level phylotypes but mostly non-overlapping populations of isolates when grouped lower than the level of genus. Further, correlations between bacterial phylotype and SM production varied at the species level and below, suggesting SM distribution within bacterial taxa must be analyzed on a case-by-case basis. Our results suggest that two *E. fragilis* freshwater sponges collected in similar environments can exhibit large culturable diversity on a species-level scale, thus researchers should scrutinize the isolates with analyses that take both phylogeny and SM production into account in order to optimize the chemical space entering into a downstream bacterial library. Depending solely on 16S rRNA gene sequencing, MALDI protein spectra, or morphological analyses will not fully inform researchers of the SM potential of recovered isolates, and this will lessen the downstream efficiency within a SM discovery program.

## Materials and methods

### Sample collection and processing

Samples of *Eunapius fragilis* were collected by citizen science divers using SCUBA. Sponge SCD18 was collected July 15, 2015 from a depth of 21 m, off of the wooden substrate of the Vickery shipwreck (−76.0191, 44.28025). Sponge SC18 was collected June 17, 2015 from a depth of 21 m, off of the wooden substrate of the Iroquois shipwreck (−76.0055, 44.28025). Samples were scraped into 50 mL sterile centrifuge tubes and shipped overnight to the laboratory where they were processed immediately. Following removal of any macro-debris, 1 cm^3^ of each sponge was rinsed five times with autoclaved lake water. The sponge sample was then placed in a sterilized mortar, 10 mL of sterilized 20% glycerol was added, and ground thoroughly with a sterilized pestle for two minutes. The resulting solution was separated into 500 μL aliquots. Heat treated aliquots were held at 54 °C for 9 minutes. Dilution series of 0, 1/10, and 1/100 were created with sterilized 20% glycerol. Each agar media type contained 28 μM cycloheximide (for media recipes, see supplemental information). For each preparation condition 50 μL sponge/glycerol solution was spread evenly over the surface. Agar/nutrient plates were sealed with Parafilm^®^ and left at room temperature in the dark; these are referred to as “diversity plates”. To capture more bacterial isolates from low-nutrient/low-dilution diversity plates, colonies were isolated over a longer period (four months) and plates were regularly checked to prevent overgrowth. A total of 8 diversity plates immediately overgrew with biofilms, while fungus was prevalent and overgrew in 39, despite including cycloheximide in the isolation media. These plates and their counterparts that were *not* overgrown in both samples were removed from the dataset to allow for direct sample comparison from a total of 61 diversity plates. See “Data and code availability” for a link to matching figures for both the full and reduced datasets.

### Sponge identification

Classic taxonomic morphological analysis was carried out at the genus and species level. A set of diagnostic macro- and micro-morphotraits was focused for diagnosis (i.e., growth form, consistency, skeletal architecture, traits and dimensions of skeletal megascleres and microscleres such as siliceous spicules, gemmuloscleres morphs, and architecture of gemmules such as resting stages [9, 10]. Morphometries were performed on gemmules and spicules of each spicular type. Samples were studied by stereomicroscope for macro-traits. For skeletal analyses representative body fragments were dissected by hand, processed by dissolution of organic matter in boiling 65% nitric acid, and rinsed in tap water. After decantation, cleaned spicules were suspended in ethanol, dropped onto slides, and glued in Eukitt with a cover slide as permanent preparations. The spicular complements were compared with recent and historical literature and with registered collections. Samples were ascribed to *Eunapius fragilis* (Leidy, 1851) (Porifera, Demospongiae, Spongillida, Spongillidae). This species is common in inland water and widespread from almost all Terrestrial Ecoregions of the World (Nearctic, Palaearctic, Neotropical, Afrotropical, Oriental, and Australian). In the Nearctic Region this species was recorded from Canada and the United States (Colorado, Connecticut, Florida, Illinois, Indiana, Iowa, Kansas, Kentucky, Louisiana, Maine, Michigan, Minnesota, Montana, Newfoundland, New Jersey, New Scotland, New York, Ohio, Pennsylvania, Texas, Wisconsin, and Wyoming).

The main morphotraits of the species are the following. Growth form encrusting, variably thick. Consistency is notably soft and fragile in both *in vivo* and dry conditions. Color is whitish to greenish. Surface is slightly hispid likely due to erected spicules. Oscules are not conspicuous *in vivo*, and are scattered in a network of subdermal canals. There is a skeletal network of monaxial spicules and collagen. Megascleres exist as smooth oxeas (160–270 × 4–15 μm). Microscleres are absent. Gemmules are subspherical (300–450 μm). Gemmuloscleres exist as straight to slightly curved strongyles to strongyloxeas (35–140 × 3–8 μm) and are irregularly tangential and embedded into the gemmular theca. The habitat consists of a wide range of lentic and lotic habitats.

### MALDI-TOF MS analysis

Isolates were purified onto high nutrient agar plates (A1) and biological replicates (three colonies of each isolate) were separately applied, as a thin smear, to a 384-spot MALDI target plate (Bruker Daltonics, Billerica, MA, USA) using a sterile toothpick. As we compared protein and SM spectra, it was important to calibrate spectra and perform manual inspections to ensure correlation wasn’t due to experimental design/batch effects. MALDI-TOF MS settings can be found in previous IDBac publications [34, 49], including in a written and video protocol [50]. A minor modification was the shift in the analysis range to 4-20 kDa as opposed to 3-20 kDa, based on recent work by Strejcek et al [39]. IDBac relies on a number of R packages including mzR [67] and MALDIquant [68, 69].

### 16S-rRNA analysis

DNA from 361 bacterial isolates was extracted using a DNeasy UltraClean microbial kit (Qiagen). The 16S rRNA gene was amplified using 27F (5^′^-CAGAGTTTGATCCTGGCT-3^′^) and 1492R (5^′^-AGGAGGTGATCCAGCCGCA-3^′^) [70] primers using polymerase chain reaction (PCR) under the following conditions: initial denaturation at 95 C for 5 min; followed by 35 cycles of denaturation at 95 °C for 15 s, annealing at 60 °C for 15 s, and extension at 72 °C for 30 s; and a final extension step at 72 °C for 2 min. PCR products were purified using a QIAquick PCR purification kit from Qiagen, and the amplicons sequenced by Sanger sequencing. Data were analyzed by Geneious V11.1.4 software and genus assigned using the SILVA Alignment, Classification and Tree (ACT) Service [55].

## Data and Code Availability

MALDI-TOF MS data were deposited in MassIVE (DOI: 10.25345/C5261K, accession: MSV000087941). The IDBac database and MALDI-TOF MS spectra of previously identified bacteria used for spectra matching and identification are available from MassIVE (DOI: 10.25345/C5261K, accession: MSV000083461).

Partial and full 16S-rRNA sequences of environmental bacteria used in this study were deposited in GenBank with the accession numbers MT596540 to MT596560. Matching figures for both the full and reduced datasets, as well as the code, data, and Docker images to reproduce all analyses and figures is available from: https://doi.org/10.5281/zenodo.5123348.

## Acknowledgements

The authors wish to acknowledge Joseph Dudiak and Ken Kozen of Clayton Dive Club for collection of the freshwater sponges. Research reported in this publication was supported by the National Institute of General Medical Sciences of the National Institutes of Health under Award Number R01GM125943 (B.T.M. and Dr. Laura M. Sanchez). The content is solely the responsibility of the authors and does not necessarily represent the official views of the National Institutes of Health. This work was also supported by NIH F31 AT010419 (C.M.C.), T32 AT007533 (M.W.M.) and National Geographic grant CP-044R-17.

## Contributions

C.M.C. and B.T.M. designed the project. C.M.C., A.H., B.T.M. designed experiments. C.M.C. developed and wrote the code and performed the analyses. C.M.C., M.W.M., J.F.-H., B.T.M. gathered collection permits and organized citizen scientist divers. R.M. and R.P. were responsible for sponge identification. C.M.C. and A.H. collected MALDI data. C.M.C., E.L., S.R. collected and analyzed 16S rRNA data. C.M.C. and B.T.M. wrote the manuscript and all authors reviewed and edited the manuscript.

