## Supplemental Information for "Relationship between bacterial phylotype and specialized metabolite production in the culturable microbiome of two freshwater sponges"

Note: Except for S1, all supplemental figures were generated using the dataset derived from the 692 isolates from matching media conditions. All figures (including matching figures using the full 851 isolates), code, and data to reproduce them are available from: DOI: 10.5281/zenodo.5123348

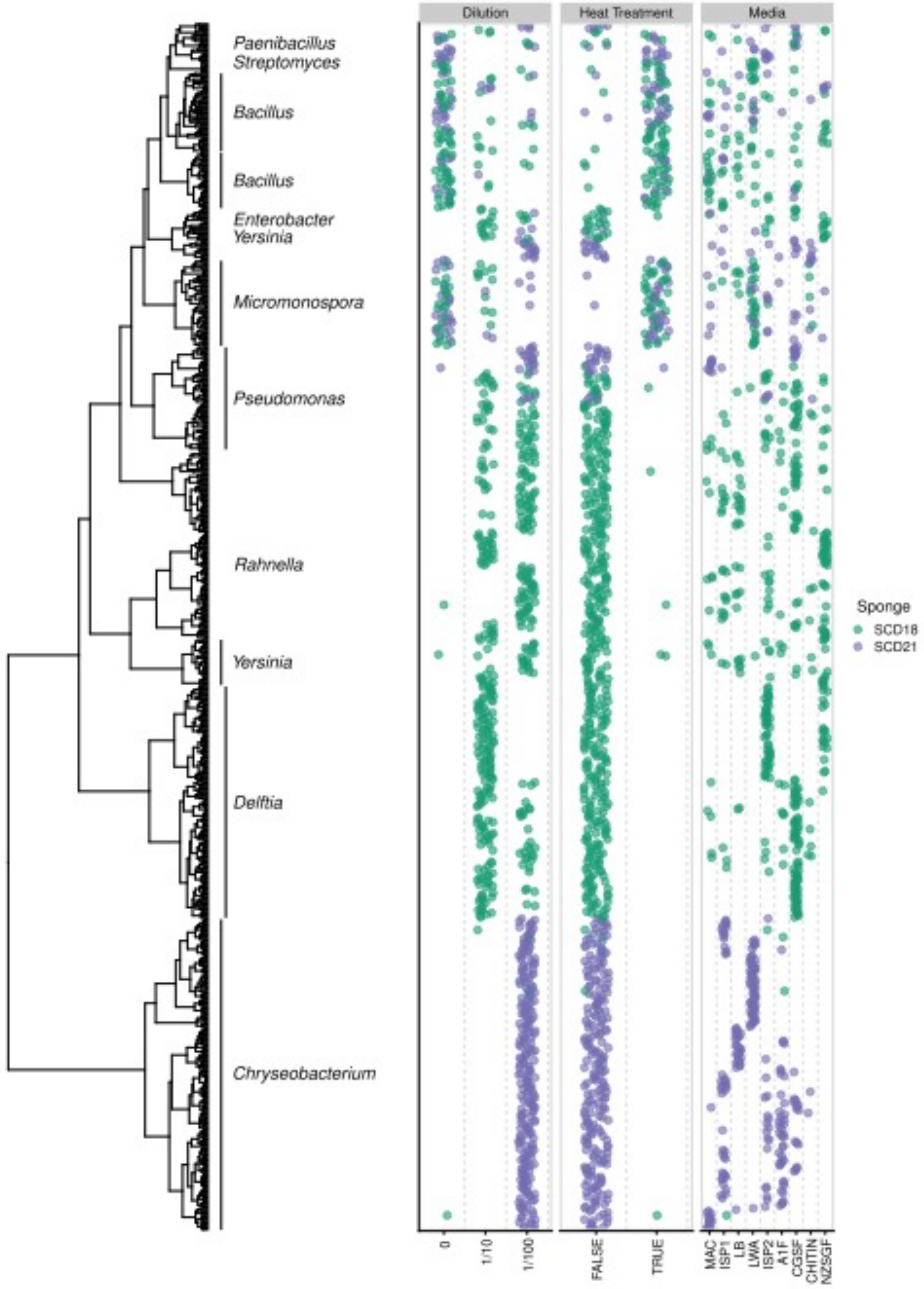


**S1: Hierarchical clustering of MALDI-TOF MS protein spectra from all 851 bacterial isolates.** Isolates identified using MALDI-TOF MS and/or 16S-rRNA gene sequence analysis were plotted beside the dendrogram along with a scatter plot displaying isolation conditions for each isolate, colored by the source sponge.

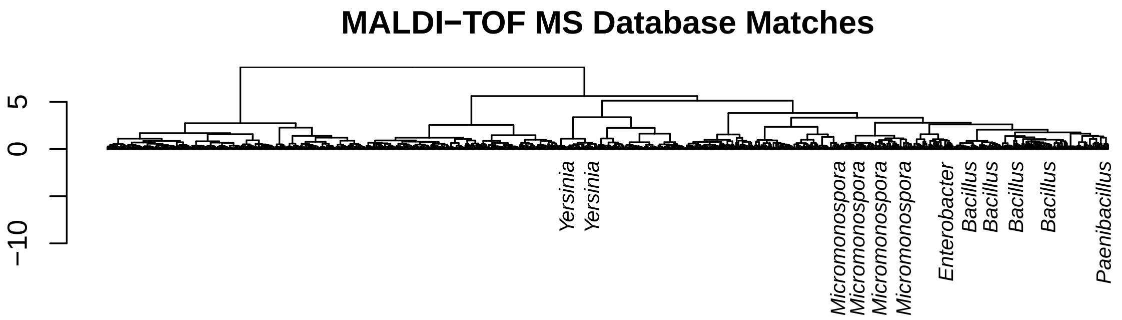


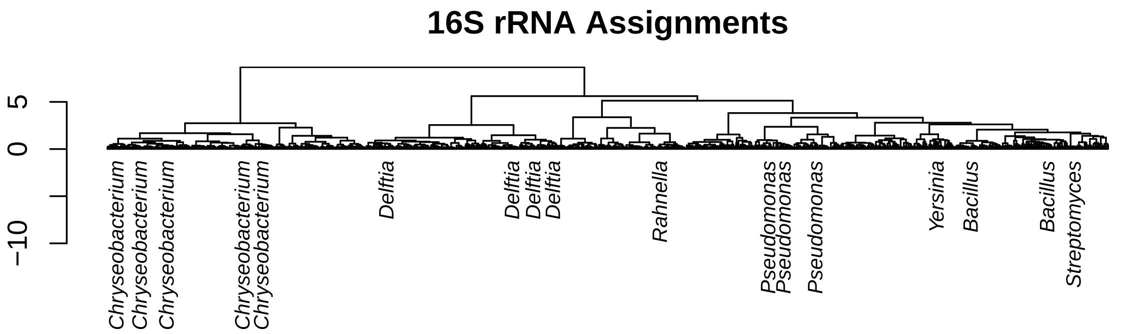


**S2: 16S rRNA gene sequence analysis and in-house database matching of MALDI-TOF MS spectra.** Both top and bottom dendrograms are identical and represent groupings based on MALDI-TOF MS analysis of the protein region (4K to 20K Da). Sponge bacterial isolates were identified using 16S rRNA gene sequencing, SILVA alignment, and database matching (see 16S rRNA sequencing section of material and methods section). We also putatively identified isolates using MALDI-TOF MS analysis, employing cosine similarity scores against a database of previously identified environmental isolates (and manual inspection of matching spectra). MALDI-TOF MS matches aligned well with the 16S rRNA assignments (*Bacillus* spp. and *Yersinia*/*Enterobacter*) and revealed additional potential *Yersinia* and *Micromonospora* isolates. Isolates within the *Micromonosopora* pseudo-phylogenetic group were morphologically congruent with *Micromonosopora* spp. Note: *Yersinia*, *Rahnella*, and *Enterobacter* all belong to the same order- Enterobacterales.

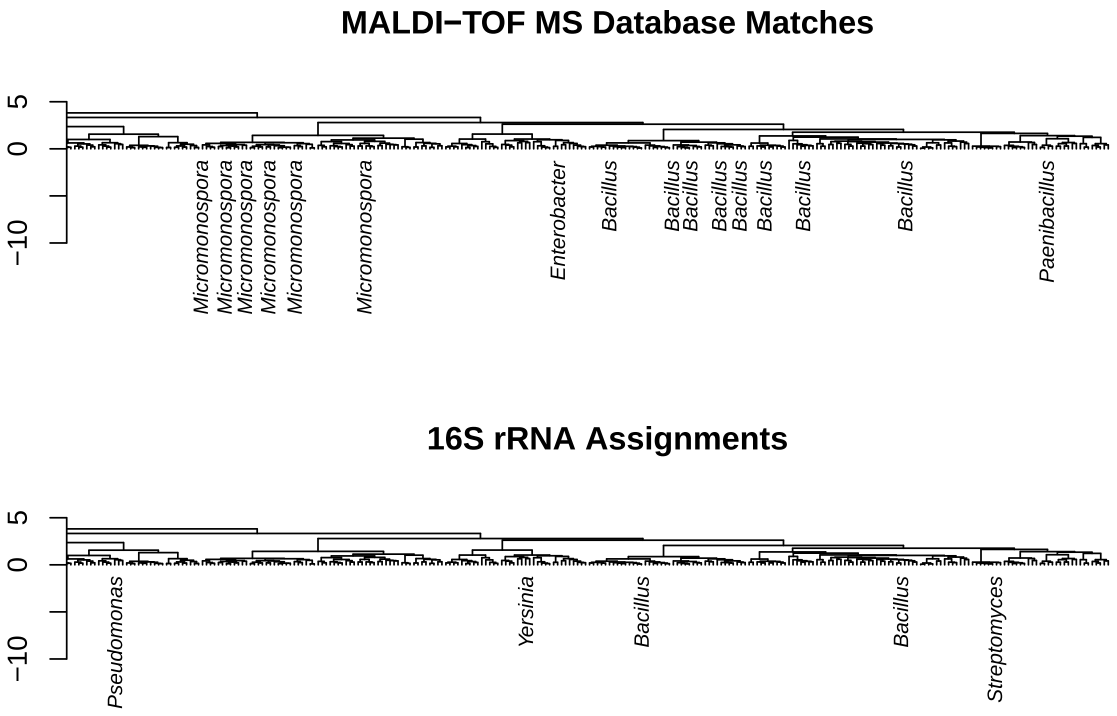


**S3: Expansion of MALDI-TOF MS protein fingerprint dendrogram highlighting congruence of MALDI-TOF MS and 16S rRNA assignment of *Bacillus* spp. and additional identification of *Micromonospora* spp. by MALDI-TOF MS.**

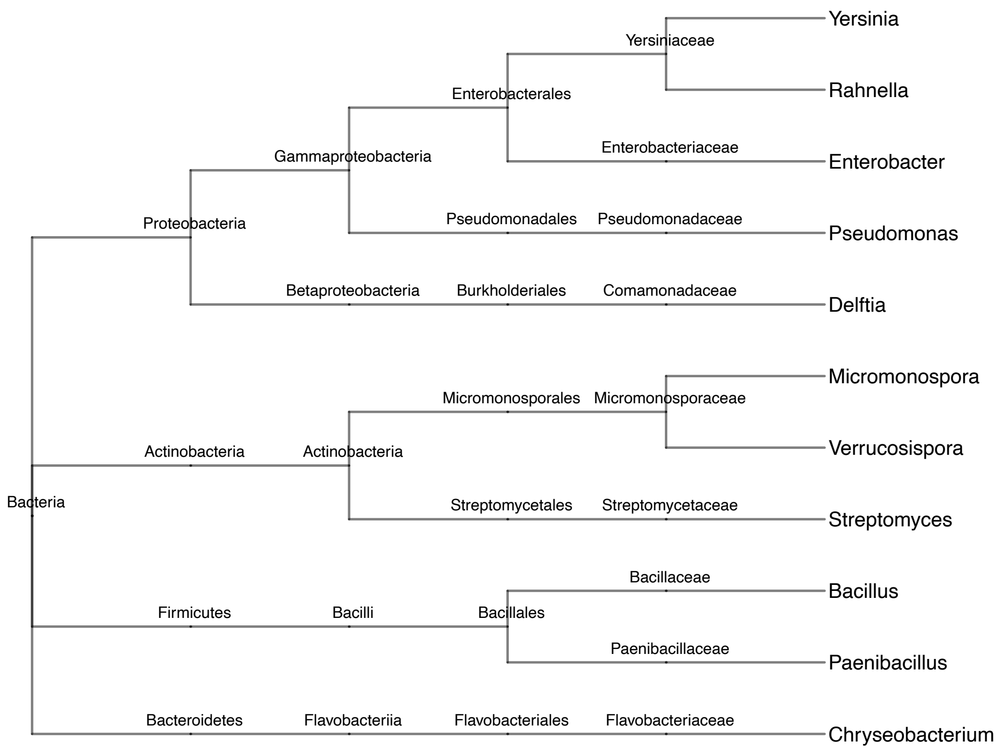


**S4: Taxonomic tree of genera recovered from *Eunapius fragilis* var *minuta*.** Created using online tools available at www.ncbi.nlm.nih.gov/Taxonomy, on 2020/05/11.

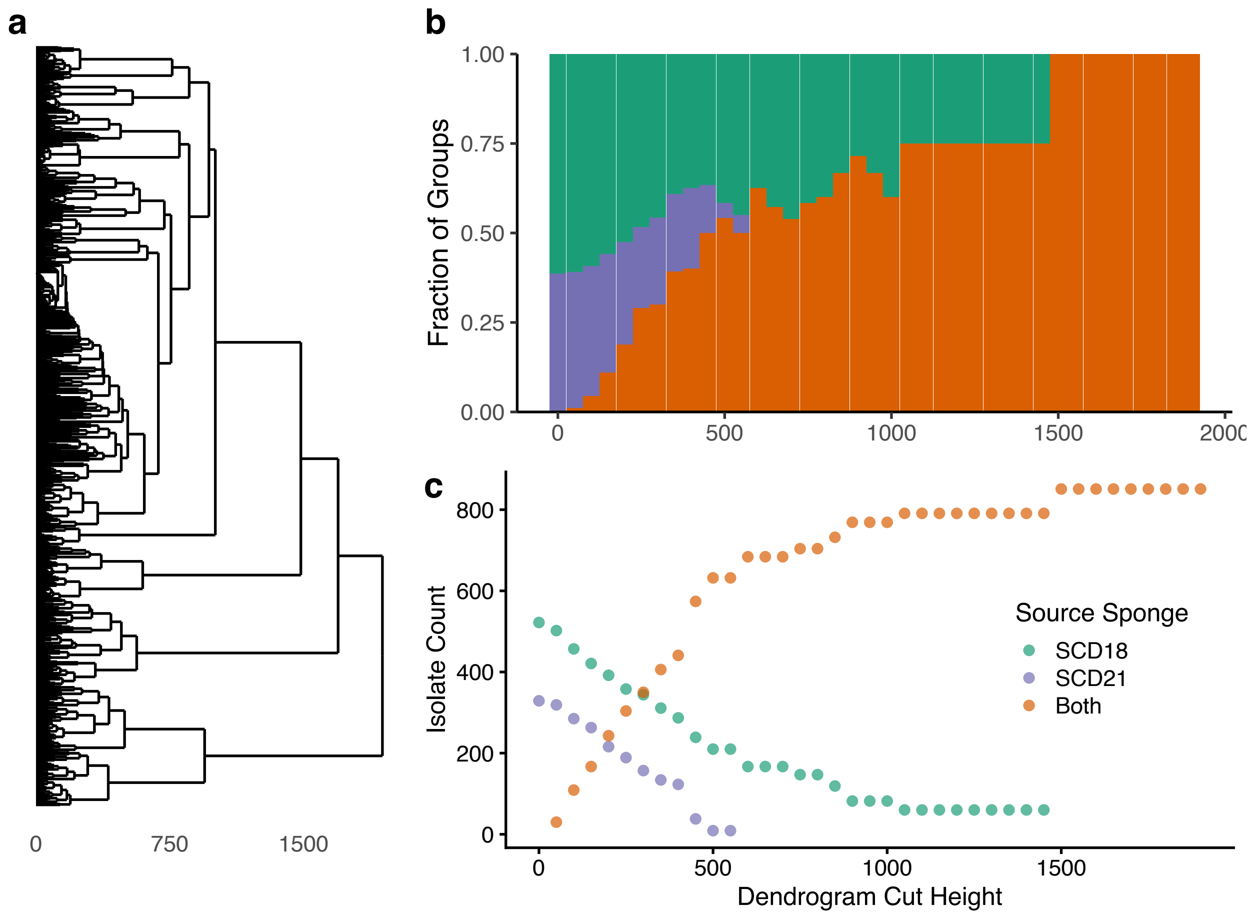


**S5:** (a) Hierarchical clustering of MALDI-TOF MS protein spectra was used to determine the overlap of isolates between sponges in a similar manner to Fig. 2 except using a different peak-binning implementation (MALDIquant rather than the feature within IDBac) and distance metric (Euclidean rather than cosine). (b) By "cutting" the dendrogram at intervals we determined the fraction of groups that contained isolates from SCD18, SCD21, or both. Because the number of groups doesn't provide insight to the number of isolates, we also evaluated (b) the raw isolate counts within groups for every cut (c). These results are consistent with the results of Figs. 2 and S6.

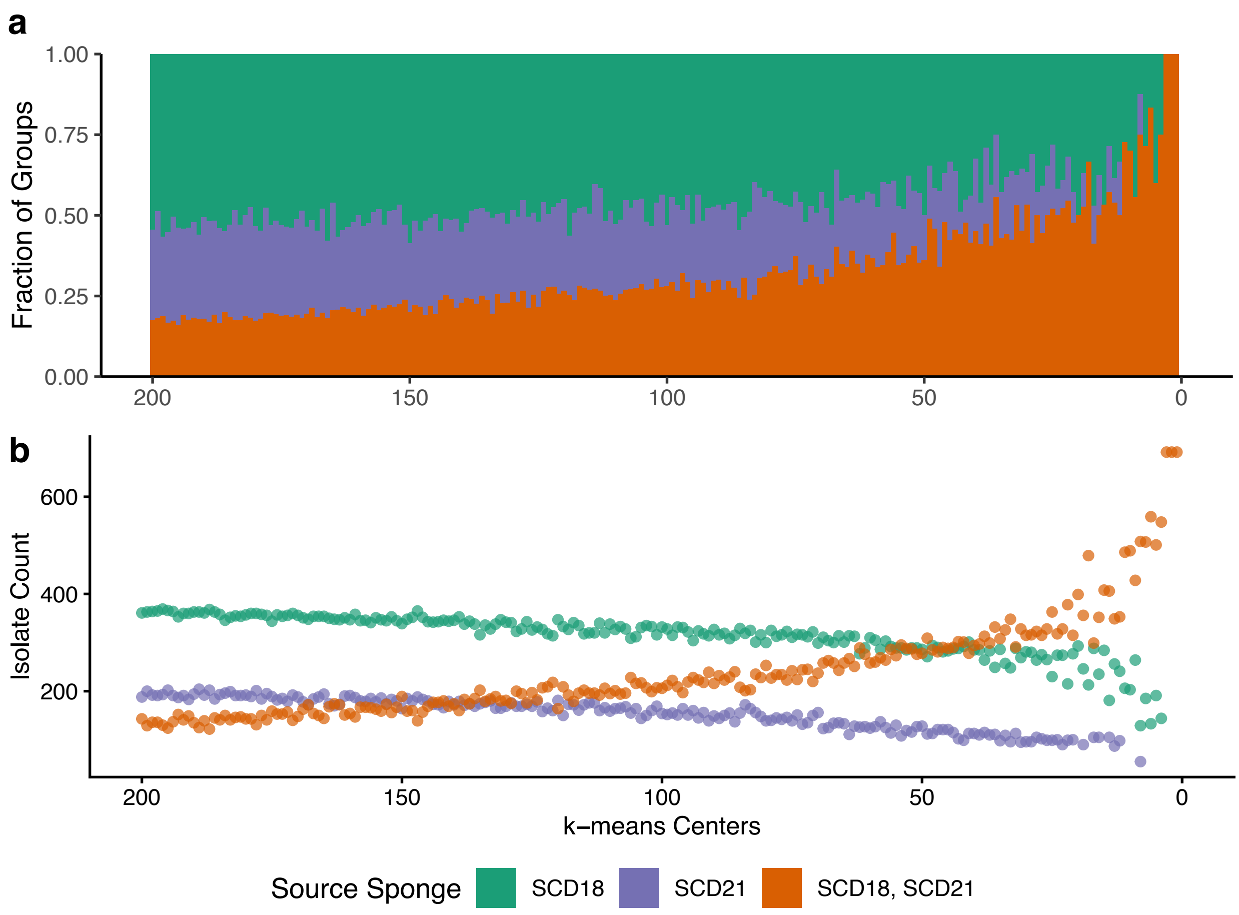


**S6:** Pseudo-phylogenetic group composition, calculated with k-means clustering. Calculating k-means, k = 1 through k = 200, we determined the fraction of groups that contained isolates from SCD18, SCD21, or both (a). We also evaluated raw isolate counts within groups for every iteration (b). These results are consistent with what was found in Fig. 2 and S5. The k-means calculations were performed with the same distance matrix used to create Figs. 1 and 2.

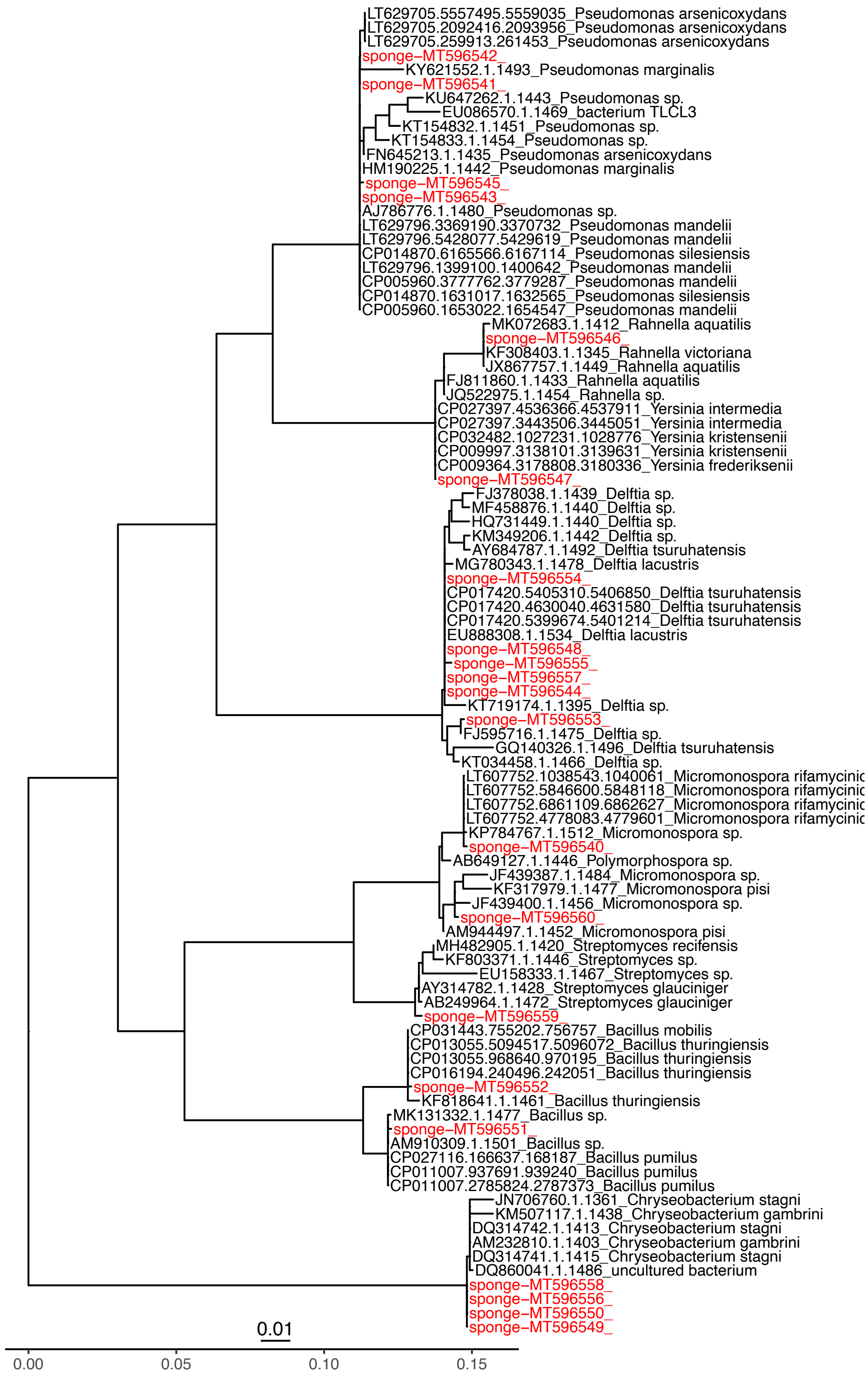


S7: caption on next page

**S7: Phylogenetic tree created from full and partial 16S-rRNA gene sequences of sponge bacterial isolates.** Red tip labels represent sponge samples and black labels represent reference strain matches. Alignments and tree creation were performed using SILVA's Alignment, Classification and Tree Service. Gene sequences were deposited in GenBank with the accession numbers MT596540 to MT596560.

Pruesse, E.; Peplies, J.; Glöckner, F. O. SINA: Accurate High-Throughput Multiple Sequence Alignment of Ribosomal RNA Genes. Bioinformatics 2012, 28 (14), 1823–1829.

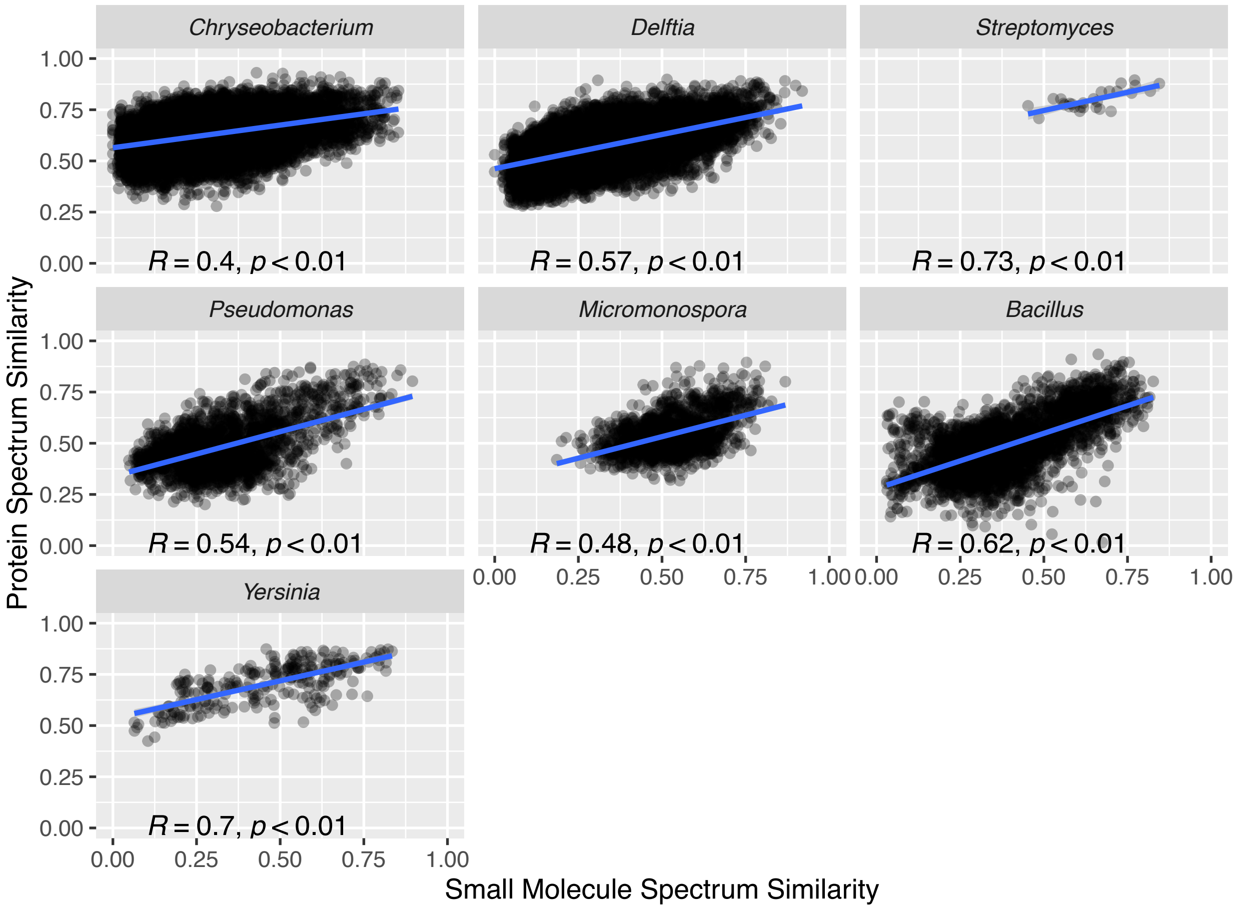


**S8: Correlation of protein and SM spectra within pseudo-phylogenetic groups.** Cosine similarity scores for protein and SM spectra were calculated pairwise within six pseudo-phylogenetic groups as defined in Fig. 1. While all genera had some positive correlation, *Bacillus* isolates showed comparatively high correlation. These results corroborate those found in the MAN analysis of Fig S11.

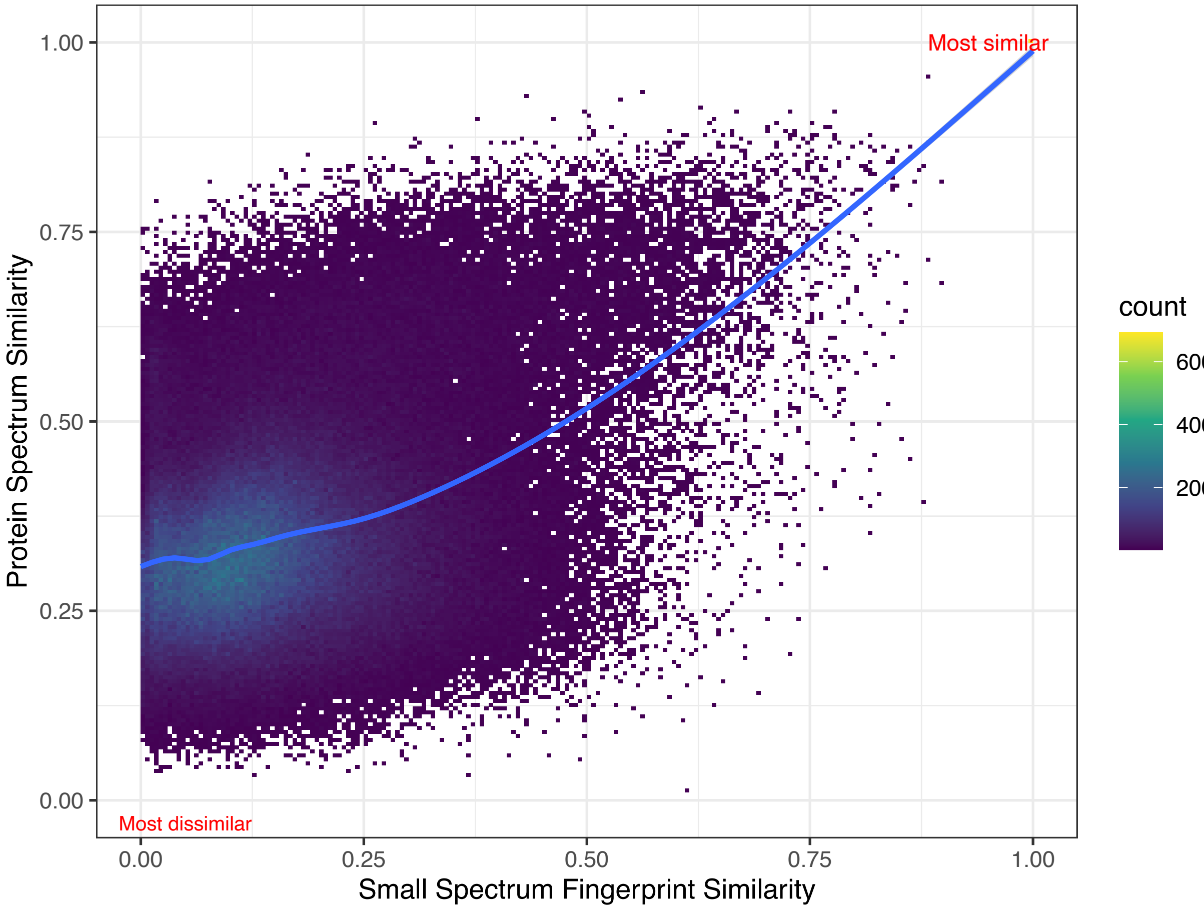


**S9: Pairwise protein and small molecule distance comparison of bacterial isolates.** Similar to Fig. S8, here protein and SM spectra were compared, pairwise for all 692 isolates from matched agar conditions. Most comparisons are low similarity for both types of spectra, as expected. However, as displayed by the blue-line (default local regression using ggplot::geom_smooth), as protein spectra similarity increases, specialized metabolite spectra similarity also increases. Data are represented as bins of observations using ggplot2's geom_bin2d function and bin size of 200. This means each single point in the graph represents 200 cosine similarity scores.

H. Wickham. ggplot2: Elegant Graphics for Data Analysis. Springer-Verlag New York, 2016.

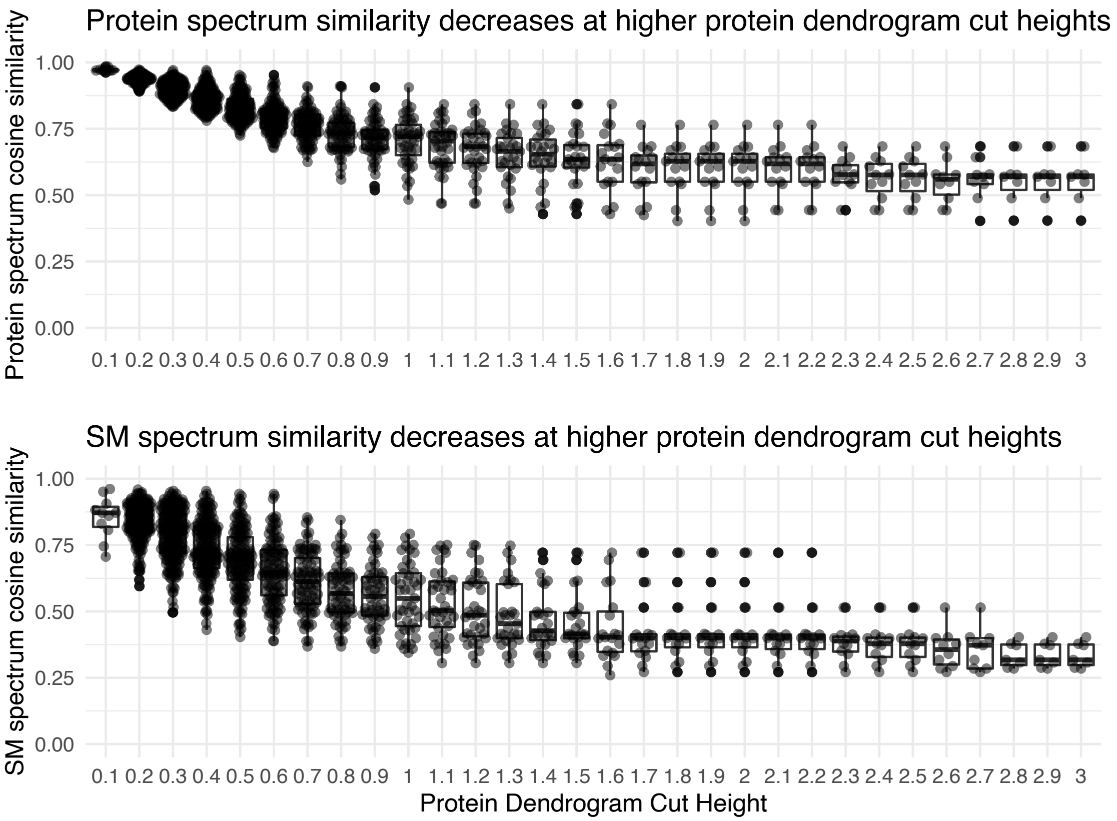


**S10: Comparing protein and specialized metabolite spectra cosine similarity.** The dendrogram described in Fig. 1 was cut/divided at 0.1 intervals from a height of 0.1 to 3.0. At each cut the median cosine similarity in each group was calculated for both protein and SM spectra. As expected, protein spectrum similarity decreases at higher dendrogram cut heights. Additionally, as seen in the lower panel, SM spectra also become less similar as pseudo-phylogenetic groups are “loosened” to contain isolates with less similar protein spectra. In other words, SM spectra were most similar when their protein spectra were also similar, supporting that, broadly speaking, SM production patterns correlate with phylogenetic groupings. Observations decrease from left to right because there are fewer groups from which to make intra-group comparisons.

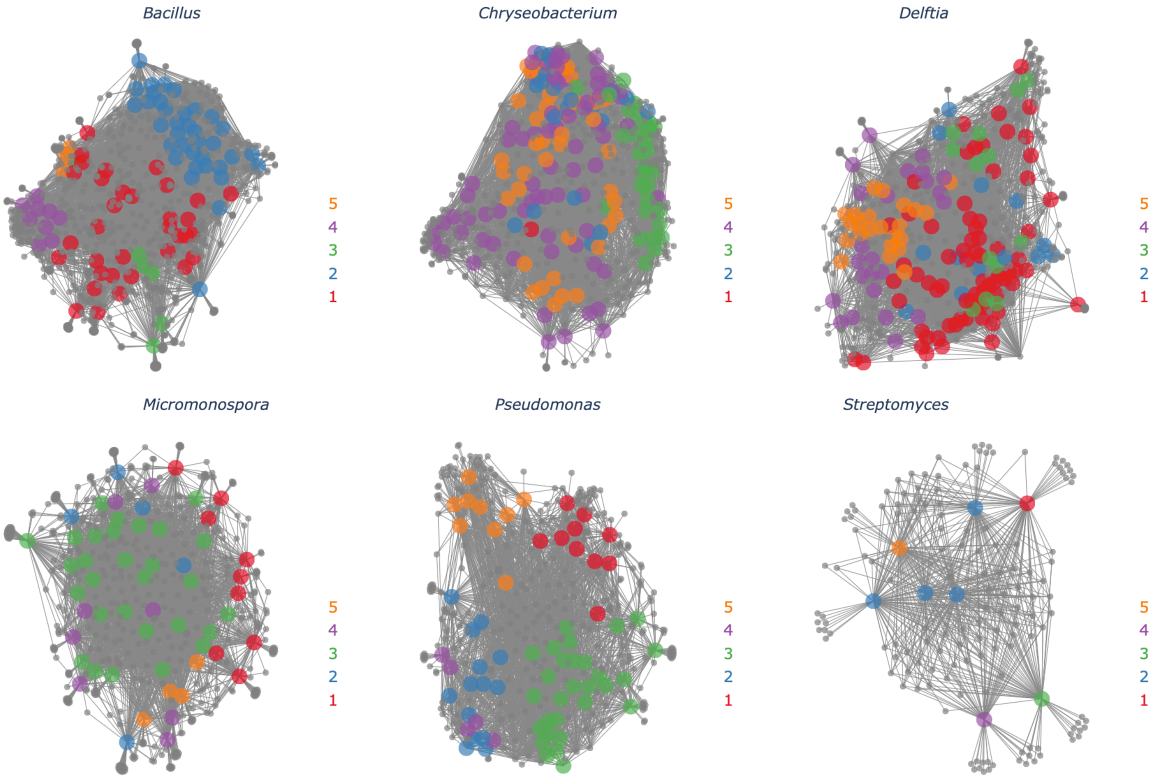


**S11a: Metabolite Association Network of freshwater sponge bacterial genera, colored by protein spectrum similarity.** Intra-genus variability in SM production is shown by splitting isolates into groups of five, determined by protein spectrum similarity (as in Fig. 3). As observed within the *Bacillus* and *Pseudomonas* MANs, subgroups often exhibited distinct SM production, corresponding directly to pseudo-phylogenetic variation.

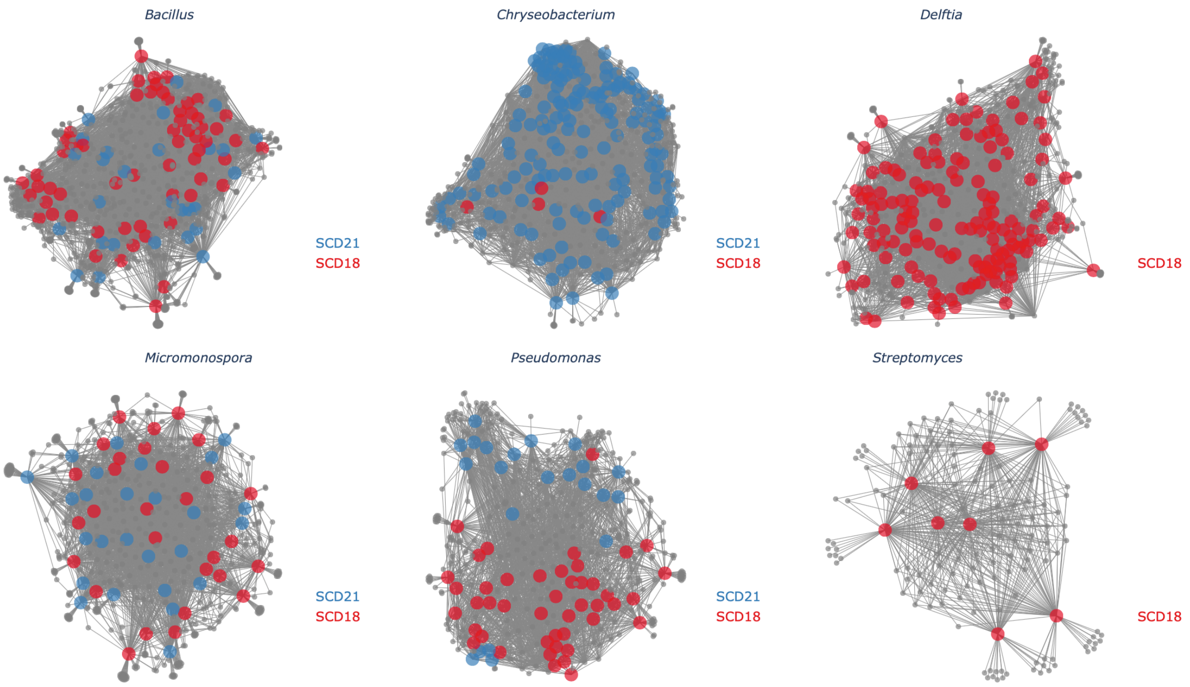


**S11b: Metabolite Association Network of freshwater sponge bacterial genera, colored by host sponge.** Here, isolate nodes are colored according to the sponge source (SCD18 or SCD21). See manuscript for discussion on the limitation of confounding.

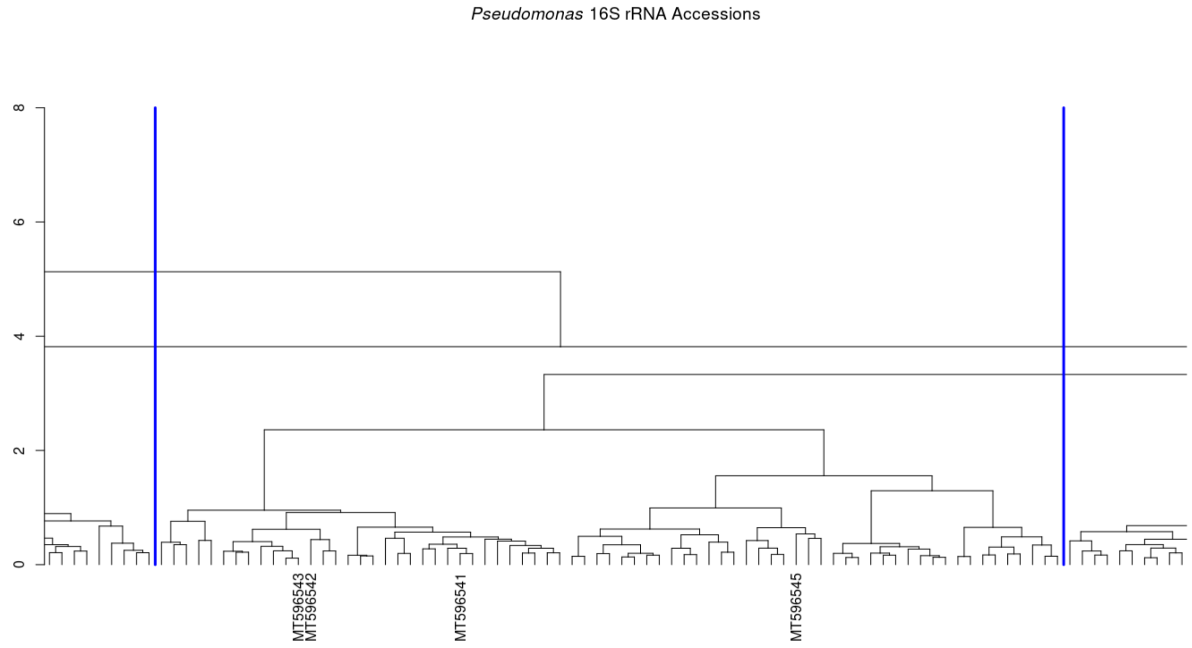


**S12: Protein dendrogram zoom-in of all isolates (Fig. S1), showing the positions of *Pseudomonas* isolates identified by 16S rRNA sequencing analysis (Fig. S7).** Blue lines denote the boundaries of isolates defined in the study as "*Pseudomonas*".

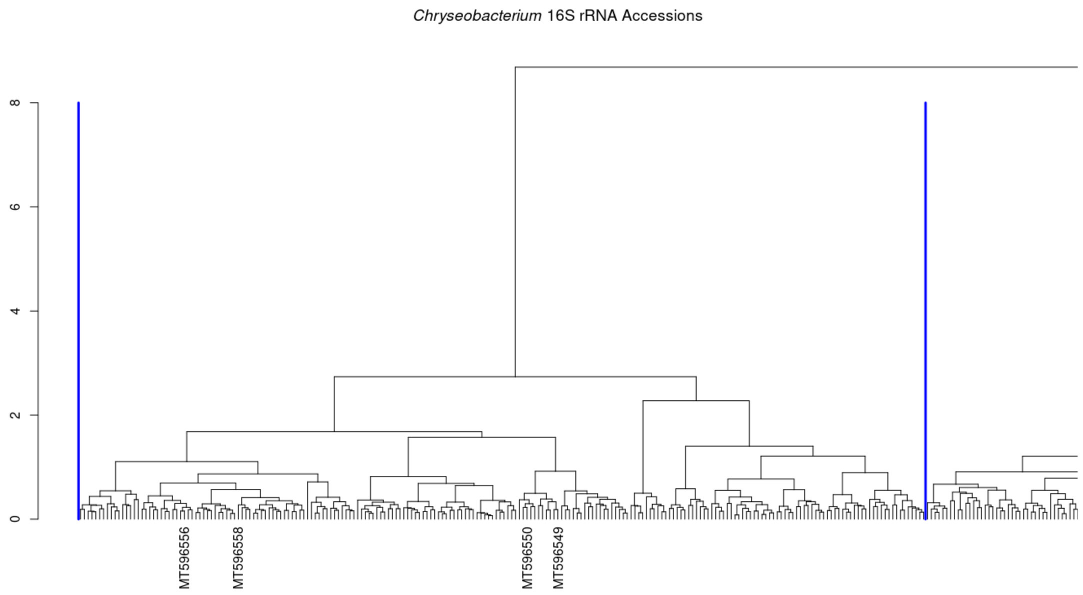


**S13: Protein dendrogram zoom-in of all isolates (Fig. S1), showing the positions of *Chryseobacterium* isolates identified by 16S rRNA sequencing analysis (Fig. S7).** Blue lines denote the boundaries of isolates defined in the study as "*Chryseobacterium*". The group of *Chryseobacterium* isolates missing 16S-rRNA sequencing to the right of the grouping were identified by MALDI MS analysis (Fig. S14).

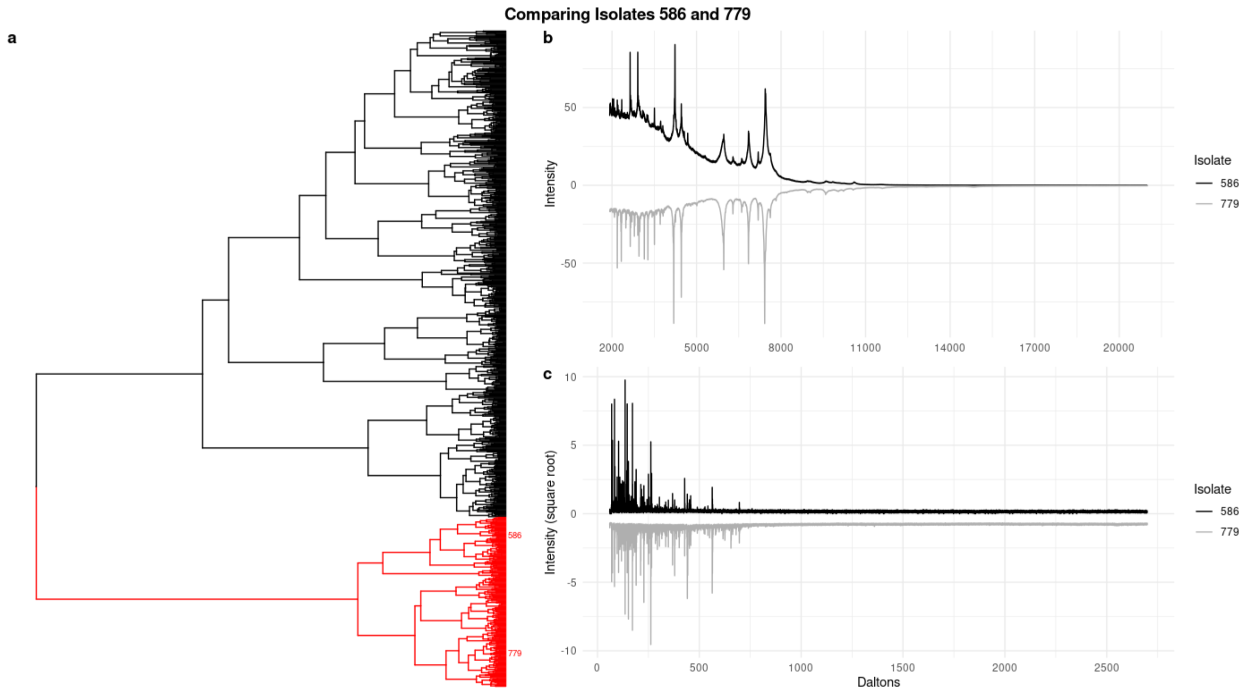


**S14: Identifying additional *Chryseobacterium* isolates through MALDI-TOF MS spectra comparisons.** All groupings in the study that were assigned a genus ID were identified by 16S rRNA or MALDI-TOF MS protein spectra comparisons (Fig. S2-3). Boundaries of the groupings were further delineated by comparing multiple spectra across close groupings using a custom R script that created multi-plot figures exactly as shown here. Provided two isolate identifiers (586 and 779 in a-c), the full data dendrogram (Fig. S1) was plotted and the sub-dendrogram representing the least common ancestor (LCA) node was colored red (a). The similarity in the mean protein spectrum was then compared and only high quality and highly similar spectra were considered as matches (b). Lastly, the specialized metabolite spectrum was also evaluated for similarity/differences (c). Here we show a representative comparison from evaluating the group of putative *Chryseobacterium* isolates without 16S rRNA coverage. As shown from the position of the isolates in the dendrogram (a) and the high similarity between protein spectra (b) we were confident in assigning the group as *Chryseobacterium*. This comparison was done multiple times using isolates from multiple areas across the *Chryseobacterium* grouping.

| Ingredients | LWA | NZSG | A1 | ISP1 | ISP2 | CHITIN | LB | CGS | MAC |
|---|---|---|---|---|---|---|---|---|---|
| Malt Extract (g) | 0.0 | 0.0 | 0.0 | 0.00 | 10.0 | 0.000 | 0.0 | 0.0 | 0.0 |
| Yeast Extract (g) | 0.0 | 5.0 | 2.0 | 1.50 | 2.0 | 0.000 | 2.5 | 0.0 | 0.0 |
| Soluble Starch (g) | 0.0 | 10.0 | 5.0 | 0.00 | 0.0 | 0.000 | 0.0 | 0.0 | 0.0 |
| Glucose (g) | 0.0 | 5.0 | 0.0 | 0.00 | 2.0 | 0.000 | 0.0 | 0.0 | 1.5 |
| 50% Glycerol (ml) | 0.0 | 0.0 | 0.0 | 0.00 | 0.0 | 0.000 | 0.0 | 10.0 | 5.0 |
| N-Z Amine A (g) | 0.0 | 2.5 | 0.0 | 0.00 | 0.0 | 0.000 | 0.0 | 0.0 | 0.0 |
| Calcium Carbonate (g) | 0.0 | 3.0 | 0.0 | 0.00 | 0.0 | 0.000 | 0.0 | 0.0 | 0.0 |
| Casamino acids (g) | 0.0 | 0.0 | 0.0 | 0.00 | 0.0 | 0.000 | 0.0 | 2.0 | 1.5 |
| Casitone (g) | 0.0 | 0.0 | 0.0 | 1.25 | 0.0 | 5.000 | 0.0 | 0.0 | 1.5 |
| Peptone (g) | 0.0 | 0.0 | 1.0 | 0.00 | 0.0 | 0.000 | 0.0 | 0.0 | 0.0 |
| Soytone (g) | 0.0 | 0.0 | 0.0 | 0.00 | 0.0 | 0.000 | 0.0 | 2.5 | 1.5 |
| Tryptone (g) | 0.0 | 0.0 | 0.0 | 0.00 | 0.0 | 0.000 | 5.0 | 0.0 | 1.5 |
| NaCl (g) | 0.0 | 0.0 | 0.0 | 0.00 | 0.0 | 0.000 | 5.0 | 0.0 | 0.0 |
| K2HPO4 (g) | 0.0 | 0.0 | 0.0 | 0.00 | 0.0 | 0.700 | 0.0 | 0.0 | 0.0 |
| KH2PO4 (g) | 0.0 | 0.0 | 0.0 | 0.00 | 0.0 | 0.300 | 0.0 | 0.0 | 0.0 |
| MgSO4-7H20 (g) | 0.0 | 0.0 | 0.0 | 0.00 | 0.0 | 0.500 | 0.0 | 0.0 | 0.0 |
| 8 mg/mL FeSO4-7H2O (ml) | 0.0 | 0.0 | 0.0 | 0.00 | 0.0 | 0.625 | 0.0 | 0.0 | 0.0 |
| Chitin (g) | 0.0 | 0.0 | 0.0 | 0.00 | 0.0 | 4.000 | 0.0 | 0.0 | 0.0 |
| Agar (g) | 7.5 | 7.5 | 7.5 | 7.50 | 7.5 | 7.500 | 7.5 | 7.5 | 7.5 |
| Filtered Lake Water (ml) | 1000.0 | 1000.0 | 1000.0 | 1000.00 | 1000.0 | 1000.000 | 1000.0 | 1000.0 | 1000.0 |

**T1: Media recipes for culturing bacteria on agar.** Freshwater sponge isolates were isolated from agar plates as described in this table (note: these are half-strength of our regular recipe plates). All MALDI experiments used isolates grown on full-strength A1.
